# Reduced oocyte quality and exacerbation of the maternal age effect are enduring consequences of low-level atrazine exposure in mouse

**DOI:** 10.1101/2022.08.23.505013

**Authors:** Yan Yun, Sunkyung Lee, Christina So, Rushali Manhas, Carol Kim, Tabitha Wibowo, Michael Hori, Neil Hunter

## Abstract

**BACKGROUND:** Egg development has unique features that render it vulnerable to environmental perturbation. The herbicide atrazine is an endocrine disruptor shown to have detrimental effects on reproduction across a number of vertebrate species.

**OBJECTIVES:** To determine whether exposure to low levels of atrazine impairs meiosis in female mammals using a mouse model; in particular, whether and how the fidelity of oocyte chromosome segregation is affected, and whether the aging-related aneuploidy is exacerbated.

**METHODS:** Female C57BL/6J mice were exposed to two levels of atrazine in drinking water, with the lower level corresponding to detected environmental contamination. To model exposure during development, atrazine was ingested by pregnant females at 0.5 days post coitum and continued until pups were weaned at 21 days post-partum. For adult exposure, 2-month-old females ingested atrazine for 3 months. For each exposure group, various indicators of oocyte quality were determined, including developmental capacity and chromosomal abnormalities during the two meiotic divisions.

**RESULTS:** Developmental exposure caused only minor effects on the fetal events of meiotic prophase-I and establishment of initial follicle pools. However, ovulation was enhanced while oocyte quality was significantly reduced. At the chromosome level, misalignment and numerical and structural abnormalities were increased at both meiotic divisions. Furthermore, fertilization efficiency was impaired *in vitro*, and apoptosis was elevated in blastocysts derived from the eggs of atrazine-exposed females. Similar levels of chromosomal defects were seen in oocytes following both developmental and adult exposure regimens suggesting that quiescent primordial follicles may be the consequential targets of atrazine. Importantly, defects were observed long after exposure was terminated. Moreover, dramatic increases in chromosomally abnormal oocytes were seen in older mice indicating that atrazine exposure during development exacerbates the effects of maternal aging on oocyte quality. Indeed, analogous to the effects of maternal age, atrazine exposure resulted in weakened cohesion between sister chromatids.

**CONCLUSION:** Low-level atrazine exposure causes persistent changes to the female mammalian germline with potential consequences for reproductive lifespan and congenital disease.

## Introduction

An egg that is competent for fertilization and development has a complex and protracted genesis that makes it uniquely vulnerable to environmental assaults (Canipari et al. 2020). Oogenesis begins during fetal development with germ-cell proliferation and differentiation into oogonia. Meiotic prophase I then ensues during which homologous chromosomes pair and undergo crossing over. Crossovers, in combination with cohesion between sister chromatids, establish connections called chiasmata that direct accurate homolog disjunction during the meiosis-I division (MI). In mammals, oocytes enter an extended arrest at the end of prophase I, coincident with the assembly of primordial follicles (in which the oocyte is surrounded by a single layer of flattened granulosa cells). Resumption of meiosis and MI occurs only in follicles selected for maturation and ovulation (Jones 2008). Thus, female fecundity is shaped by the ability of arrested oocytes to maintain their integrity over a long lifespan (up to 50 years in humans). For these and other reasons, mammalian oocytes are innately prone to chromosome segregation errors, with particularly high levels occurring in humans (Gruhn et al. 2019; Nagaoka et al. 2011). Moreover, segregation errors increase dramatically with advancing maternal age, due in large part to declining sister-chromatid cohesion (Chiang et al. 2010; Duncan et al. 2012; Gruhn et al. 2019; Jessberger 2012; Lagirand-Cantaloube et al. 2017; Lister et al. 2010).

Oocyte number and quality can be impacted by a variety of intrinsic and extrinsic factors, including exposure to environmental contaminants. The herbicide atrazine (2-chloro-4-ethylamino-6-isopropylamino-*s*-triazine) is notable as a ubiquitous environmental contaminant and potent endocrine disruptor (Hayes et al. 2011). First registered as an herbicide in the1950s, atrazine is applied to agricultural and recreational lands worldwide. The slow degradation of atrazine makes it a persistent contaminant of soil, surface and ground water, and a frequent contaminant detected in drinking water (He et al. 2019). The United States Environmental Protection Agency has defined ≤ 3 μg/L as the safe limit in drinking water supplies, while the World Health Organization has set a much higher level of 100 μg/L. Surface water contamination detected in agricultural areas is generally below 2 μg/L, but levels over 100 μg/L have been reported at some sites in the U.S. (Hatfield et al. 1996; Murphy et al. 2006). Atrazine exposure has been associated with neurotoxicity, cancer and reproductive defects (Alavanja et al. 2004; Wirbisky and Freeman 2015). Acting as an endocrine disruptor, environmental levels of atrazine cause defective steroidogenesis, gonadal dysgenesis and hermaphroditism in amphibian and fish models (Hayes et al. 2002; Rohr and McCoy 2010; Wirbisky and Freeman 2015).

In mammalian models, gestational atrazine exposure of male rats can alter hormone levels and delay puberty, although results have been somewhat conflicting. One study reported an increase in serum testosterone at 120 days post-partum (dpp) (Stanko et al. 2010), while another showed a decrease at 60 dpp (Rosenberg et al. 2008). Adult exposure in male rats is associated with reduced testosterone, delayed meiosis, and reductions in the number and quality of sperm (Abarikwu et al. 2010; Song et al. 2014). Ominously, in mouse, atrazine caused heritable epigenetic changes in the male germline (Gely-Pernot et al. 2015). Importantly, an association between atrazine exposure and poor semen quality has been established in humans (Swan et al. 2003). In females, acute exposure to high levels of atrazine during gestation can perturb oocyte meiotic prophase I and the development of primordial follicles (Gely-Pernot et al. 2017), and delay puberty (Davis et al. 2011). Exposure during adulthood was associated with increased levels of progesterone (Goldman et al. 2013), altered estrus cycles and increased atretic follicles in rats (Shibayama et al. 2009; Taketa et al. 2011). However, the effects of atrazine exposure on meiotic divisions and the fidelity of chromosome segregation have not been analyzed.

An important caveat to many mammalian studies is the high exposure doses that are generally employed; levels that are unlikely to be encountered in environment (Gely-Pernot et al. 2017; Rayner et al. 2005; Taketa et al. 2011). The exceptions detected increased progesterone, decreased oestradiol-17 beta and altered estrus in sows that were exposed to just 1 or 2 mg/kg/day in feed for 19 days, raising concerns for human exposure and reproductive health (Gojmerac et al. 1996, 1999). Indeed, epidemiological studies suggest that atrazine exposure is associated with fetal growth retardation and increased birth defects, though conflicting data exist (Chevrier et al. 2011; Munger et al. 1997; Stayner et al. 2017; Villanueva et al. 2005; Wirbisky and Freeman 2015).

Whether environmentally relevant atrazine exposure impairs female reproduction and the specific processes that are impacted remain unclear. In this study, we chose female mice as an exposure model since methods to study meiotic chromosome metabolism in mouse oocytes are well established, and the fundamental processes are conserved in humans. By exposing mice to atrazine via drinking water we addressed: (i) whether environmentally relevant levels of atrazine impact oocyte quality; (ii) the specific aspects of meiosis affected by atrazine, with a focus on chromosomal errors; (iii) whether exposure during development or adulthood have distinct effects; (iv) whether there are long-term effects on oocyte quality after exposure is terminated; and (v) whether atrazine exposure during development compounds the effects of maternal age on oocyte quality.

## Materials and Methods

### Animals and Atrazine Exposure

Founder C57BL/6J mice (Stock No: 000664) were purchased from the Jackson Laboratory. Animals were maintained in a local vivarium with temperature control and a 12-hour light/dark cycle. Weanling females (21 days post-partum; dpp) were housed, up to 4 per cage, with *ad libitum* access to standard rodent chow and water. All animals were used for experimentation according to the guidelines of the Institutional Animal Care and Use Committees of the University of California, Davis (protocol #21613). Atrazine (Molecular Formula: C8H14ClN5; Part #: N-11106-250MG; CAS: 1912-24-9; Purity: 99.3%) was purchased from Chem Service, Inc. For developmental exposure, 33 mg/L (high dose) or 100 µg/L (low dose) atrazine-containing drinking water was supplied to 2-month old pregnant females at embryonic day 0.5 (E0.5) until either euthanasia at E18.5 or until pups were weaned at 21 ddp. Weanling females were housed up to 4 per cage and euthanized for oocyte analysis by carbon dioxide overdose at 3-months old unless noted otherwise. For adult ovary exposure, 2-month-old females were supplied with high or low dose atrazine-containing drinking water for 3 months prior to euthanasia and oocyte analysis. 33 mg/L was selected as the high dose because atrazine reaches saturation at this level in water at 25 °C (Yalkowsky et al. 2019). 100 µg/L was selected as the low dose because this is the level set as the safe limit in drinking water by the World Health Organization and has been detected in contaminated water at some sites in the U.S (Hatfield et al. 1996; Murphy et al. 2006). Littermates or age-matched females were allocated to treatment groups randomly.

### Oocyte Collection and *In Vitro* Maturation

Germinal-vesicle (GV) stage oocytes were retrieved from the dissected ovaries of experimental animals without prior hormonal stimulation. Only oocytes with integral cumulus cell layers were utilized. Oocytes were collected in M2 medium (Sigma-Aldrich, M7167) under mineral oil (Sigma-Aldrich, M8410 and Nidacon, NO-100) at 37°C and cultured for *in vitro* maturation after mechanically removing surrounding cumulus cells for observation of germinal vesicle breakdown (GVBD) and polar body extrusion (PBE). GVBD was scored after 3 hrs of culture. Only oocytes that underwent GVBD within 3 hours were used to quantify PBE after a further 13 hrs of incubation.

### Chromosome Preparations From Metaphase Oocytes and Immunofluorescence

Metaphase I and II oocytes were acquired after 7 hrs and 16 hrs of culture in M2 medium at 37°C, respectively. Metaphase chromosomes were prepared as previously described (Yun et al. 2021). Briefly, Acid Tyrode’s solution (Sigma-Aldrich, M1788) was applied to remove zona pellucida, and zona-free oocytes were spread on glass slides in 1% paraformaldehyde (Electron Microscopy Science, 19208) with 0.15% Triton-X-100 (Sigma-Aldrich, X100) and 3 mM DTT (USBiological, D8070) in H_2_O pH9.2. To improve the accuracy of chromosome counting, a single oocyte was prepared in each well of a 12-well slide (Electron Microscopy Science, 63425-05). Chromosome preparations were air dried at room temperature overnight.

All of the following procedures were performed at room temperature. Metaphase I and II oocyte chromosome preparations were incubated in blocking solution (10% normal goat serum, 3% BSA, 0.05% Triton X-100, 0.05% Sodium azide in TBS) for 1-2 hrs, followed by an overnight incubation with primary antibodies: human anti-centromere antibody (ACA or CREST; ImmunoVision, HCT-0100; 1:1000 dilution) and rabbit anti-REC8 antibody (a kind gift from Dr. Scott Keeney; 1:100 dilution). After three washes in blocking solution, slides were incubated for 1 hr with goat anti-human 555 (Thermo Fisher Scientific, A-21433; 1:1000 dilution) and goat anti-rabbit 488 (Thermo Fisher Scientific, A-11034; 1:1000) secondary antibodies. Chromosomes were stained with 4′,6-diamidino-2-phenylindole at a concentration of 5 μg/ml (DAPI; Sigma-Aldrich, D8417).

### Surface-Spread Preparations of Fetal Oocyte Nuclei and Immunofluorescence

Fetal ovaries were dissected from E18.5 female fetuses and processed for surface spreading of prophase-I oocyte nuclei as described (Yun et al. 2021). Specifically, ovaries were incubated in hypotonic extraction buffer (50 mM sucrose, 30 mM Tris–HCl pH 8.0, 17 mM trisodium citrate, 5 mM EDTA, 0.5 mM DDT, 0.5 mM PMSF) on ice for 20 mins, and then minced in a 20 µl drop of 0.1 M sucrose (Sigma-Aldrich, S0389). The cell suspension was spread on a slide (Fisher Scientific, 12-544-7) coated in 1% paraformaldehyde with 0.15% Triton-X-100 in H_2_O pH9.2 and air-dried overnight in a humid chamber. Finally, the slides were washed in 0.4% Photo Flo (Kodak, 1464510) and dried again at room temperature.

Immunofluorescence staining was performed using the following primary antibodies with incubation overnight at room temperature after blocking at room temperature for 1 hour: rabbit anti-SYCP3 (Santa Cruz, sc-33195; 1:200), mouse anti-MLH1 (Cell Signaling Technology, 3515; 1:25), human anti-centromere antibodies (ACA or CREST; ImmunoVision, HCT-0100; 1:1000). Slides were subsequently incubated with the following goat secondary antibodies for 1 hr at room temperature: anti-rabbit 568 (Thermo Fisher Scientific, A11036; 1:1000), anti-mouse 488 (Thermo Fisher Scientific, A11029; 1:1000), and anti-human DyLight 649 (Jackson Labs, 109-495-088; 1:200). Coverslips were mounted with ProLong Diamond antifade reagent (Thermo Fisher Scientific, P36970) prior to imaging.

### Spindle Staining of Metaphase I and II Oocytes

For analysis of chromosome alignment, metaphase I and II oocytes were acquired after 9 hours and 16 hours of culture in M2 medium at 37°C, respectively. Intact oocytes were fixed and permeabilized in 2% paraformaldehyde in PHEM buffer (60 mM PIPES, 25 mM HEPES, 25 mM EGTA, 4 mM MgSO4) with 0.5% Triton X-100. Blocking was performed in 7% normal goat serum (Thermo Fisher Scientific, 10000C) in PBS with 0.1% Tween-20 for overnight. Primary mouse anti-α-tubulin monoclonal antibody (Thermo Fisher Scientific, A11126; diluted 1:400 in PBS with 3% BSA, 0.1% Tween-20) was then added and incubated overnight at 4°C. Secondary antibody was goat anti-mouse 488 (Thermo Fisher Scientific, A11029; 1:1000), incubated for 1 hr at room temperature. Chromosomes were stained with Hoechst (Thermo Fisher Scientific, H3569; 20 μg/ml) prior to mounting in ProLong Diamond antifade reagent.

### In Vitro Fertilization and TUNEL Assay on Blastocysts

Adult females were hormonally primed with intraperitoneal injection of 5.0 IU pregnant mares’ serum gonadotrophin (Sigma-Aldrich, G4877-2000IU), followed by injection of 5.0 IU human chorionic gonadotropin (hCG; Sigma-Aldrich, CG10) 48 hours later. At 13-14 hours after hCG injection, metaphase II eggs were retrieved from oviducts, transferred to Human Tubal Fluid (Millipore Sigma, MR-070-D) and immediately incubated with freshly prepared sperms at a final concentration of 1-2 × 10^6^ per mL. Sperm were freshly isolated from cauda epididymides of a control male (2∼5-months old) and incubated in CARD FERTIUP Preincubation Medium (Cosmo Bio, KYD-002-EX) for 60 mins prior to use. 4 hrs after eggs-sperm incubation, sperm and cumulus cells were mechanically removed by repeated pipetting and eggs/zygotes were transferred to KSOM medium (Millipore Sigma, MR-106-D) for further culture. After 24-hrs, 2-cell stage embryos were scored as a readout of fertilization efficiency. 4-cell and blastocyte stage embryos were subsequently scored at 48 and 108 hrs respectively.

Blastocytes were fixed and permeabilized in 4% paraformaldehyde in PBS with 0.5% Triton X-100. After washing in PBS, blastocytes were subject to the TUNEL reaction using the In-Situ Cell Death Detection Kit (Roche, 11684809910) according to the manufacturer’s instructions. Hoechst (20 μg/ml) was added to fluorescently stain nuclei.

### Image Acquisition and Analysis

For prophase-I and metaphase oocyte chromosomes preparations, images were acquired using a Zeiss AxioPlan II microscope equipped with a 63X oil immersion objective and Hamamatsu ORCA-ER CCD camera. Image processing and analysis were performed using Volocity (Perkin Elmer) and ImageJ (NIH) software.

For immunostaining of intact metaphase oocytes and blastocysts, 3-dimensional imaging was performed using a Zeiss Airyscan LSM 800 confocal equipped with a 63X oil immersion objective and Axiocam camera. Spindles and chromosomes in metaphase oocytes were imaged with a z-resolution of 2.0 µm, while blastocysts were captured every 5.0 µm until the entire oocyte or embryo was covered. ImageJ (NIH) software was used for image processing and analysis.

To quantify fluorescence intensities of REC8 immunostaining, metaphase-I chromosomes from the three experimental groups were prepared and processed in parallel, and images were acquired on the same day using identical exposure settings. Individual chromosomes were selected manually using DAPI staining, and the associated mean gray values were measured for REC8 and CREST channels, subtracting background signals. Relative chromosomal REC8 intensities were calculated as a ratio of REC8 to CREST, and an average for all chromosomes in each individual metaphase-I oocyte was calculated to generate the data points shown in Figure 5F. Three independent experiments were pooled by normalizing to the mean values for oocytes from each control. For inter-kinetochore distances in metaphase-II chromosomes (Figure 5B), distances between the centers of the two CREST foci within a pair of sister chromatids were measured, and then the average from all measurements in an individual metaphase-II egg was calculated to generate the data points shown in Figure 5C.

**Figure 1.**
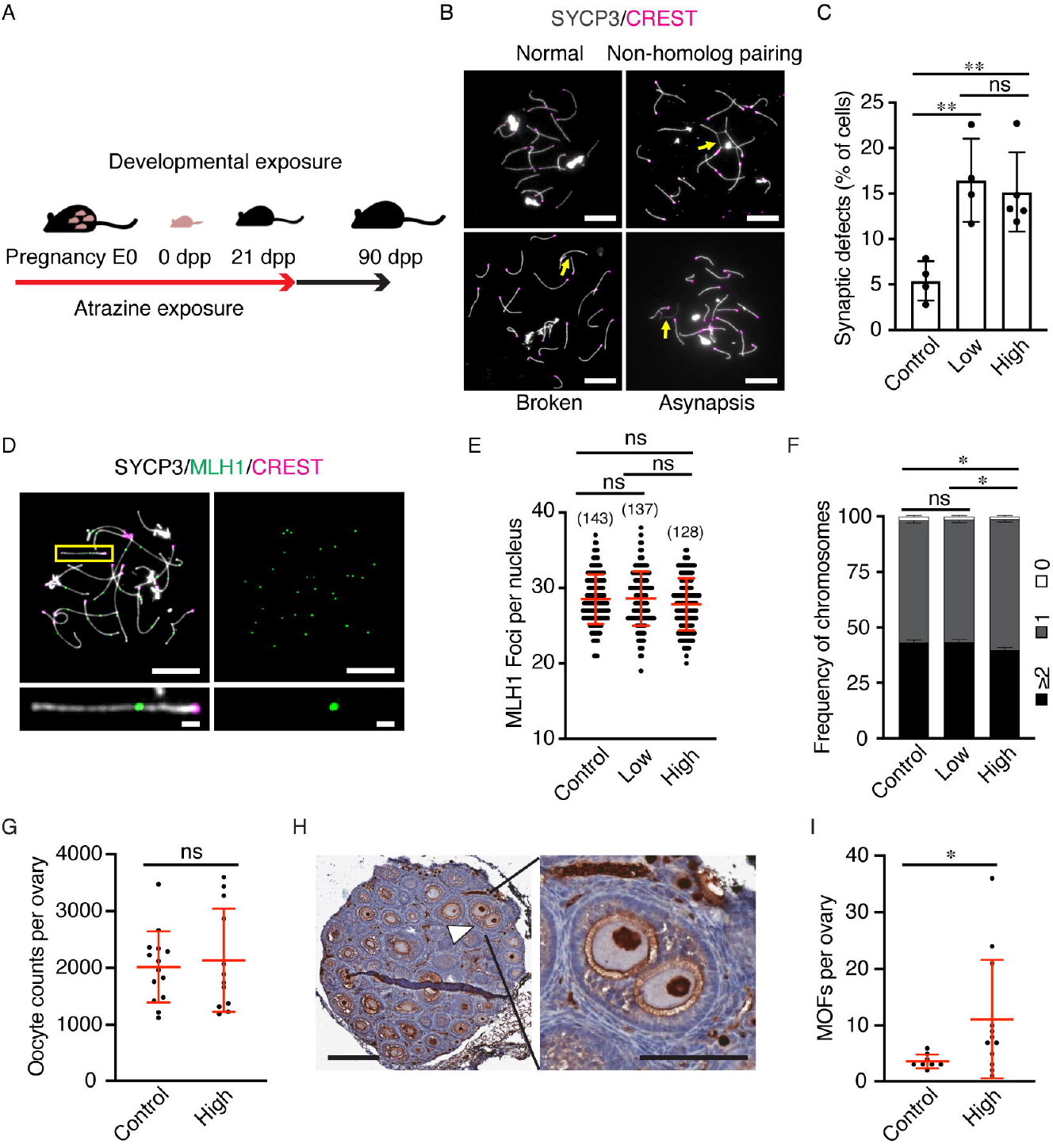
Effects of atrazine exposure during development on meiotic prophase I, ovarian reserves and multi-oocyte follicles. (*A*) Schematic of developmental atrazine exposure regimen. E0, embryonic day zero; dpp, days post-partum. (*B*) Representative images of prophase-I oocyte chromosome preparations from E18.5 embryos immunostained for SYCP3 (grey), to label homolog axes, and CREST (magenta), to label centromeres. Examples of pachytene-stage nuclei with normal synapsis, non-homologous pairing, broken axes and asynapsis are shown. Yellow arrows indicate synaptic defects. Scale bars represent 10 µm. (*C*) Quantification reveals significant increases in synaptic defects in both low and high dose atrazine groups (numbers of animals and litters used to generate data: control, 6 animals and 4 litters; low dose, 6 and 4; high dose 9 and 5). (*D*) Representative pachytene-stage oocyte chromosomes immunostained for SYCP3 (grey), CREST (magenta) and the crossover marker MLH1 (green). A single synapsed chromosome pair is magnified in the bottom panels. Scale bars represent 10 µm (main panels) and 1 µm (magnified panels). (*E*) Quantification of MLH1 foci per nucleus indicates that overall crossover numbers are not significantly changed following atrazine exposure. (*F*) Distributions of MLH1 focus numbers per chromosome show no change in the low-dose group, but a slight reduction in chromosomes with ≥2 MLH1 foci following high-dose atrazine exposure. Non-exchange chromosomes, without a MLH1 focus, were not significantly changed. Percentages of chromosomes with 0, 1, and ≥2 MLH1 foci are plotted for the three exposure groups (2860, 2740, and 2560 chromosomes in control, low and high groups, respectively; numbers of animals and litters used to generate the data in *E* and *F*: control, 10 and 4; low, 8 and 4; high 6 and 3). (*G*) Total oocyte counts per ovary from 18 dpp females (numbers of animals and litters used to generate the data: control, 7 and 6; high 6 and 2). (*H*) Representative ovary section from an 18 dpp female exposed to a high dose of atrazine, immunostained for p63 to mark oocyte nuclei and counterstained with hematoxylin. The white caret indicates a multi-oocyte follicle (MOF) containing two oocytes that is magnified in the right panel. Scale bars represent 200 µm (left panel) and 100 µm (right panel). (*I*) Numbers of MOFs per ovary (numbers of animals and litters used to generate the data: control, 4 and 2; high 6 and 2). Bidirectional error bars in *C, E, G*, and *I* represent standard deviation; unidirectional error bars in *F* represent standard error of a proportion. Data were analyzed with unpaired *t* tests (*C* and *G*) and Mann Whitney tests (*E* and *I*). The Chi-squared test was applied in *F* to compare distributions of MLH1 focus numbers. * indicates *p* < 0.05; **, *p* < 0.01; ns, not significant.

**Figure 2.**
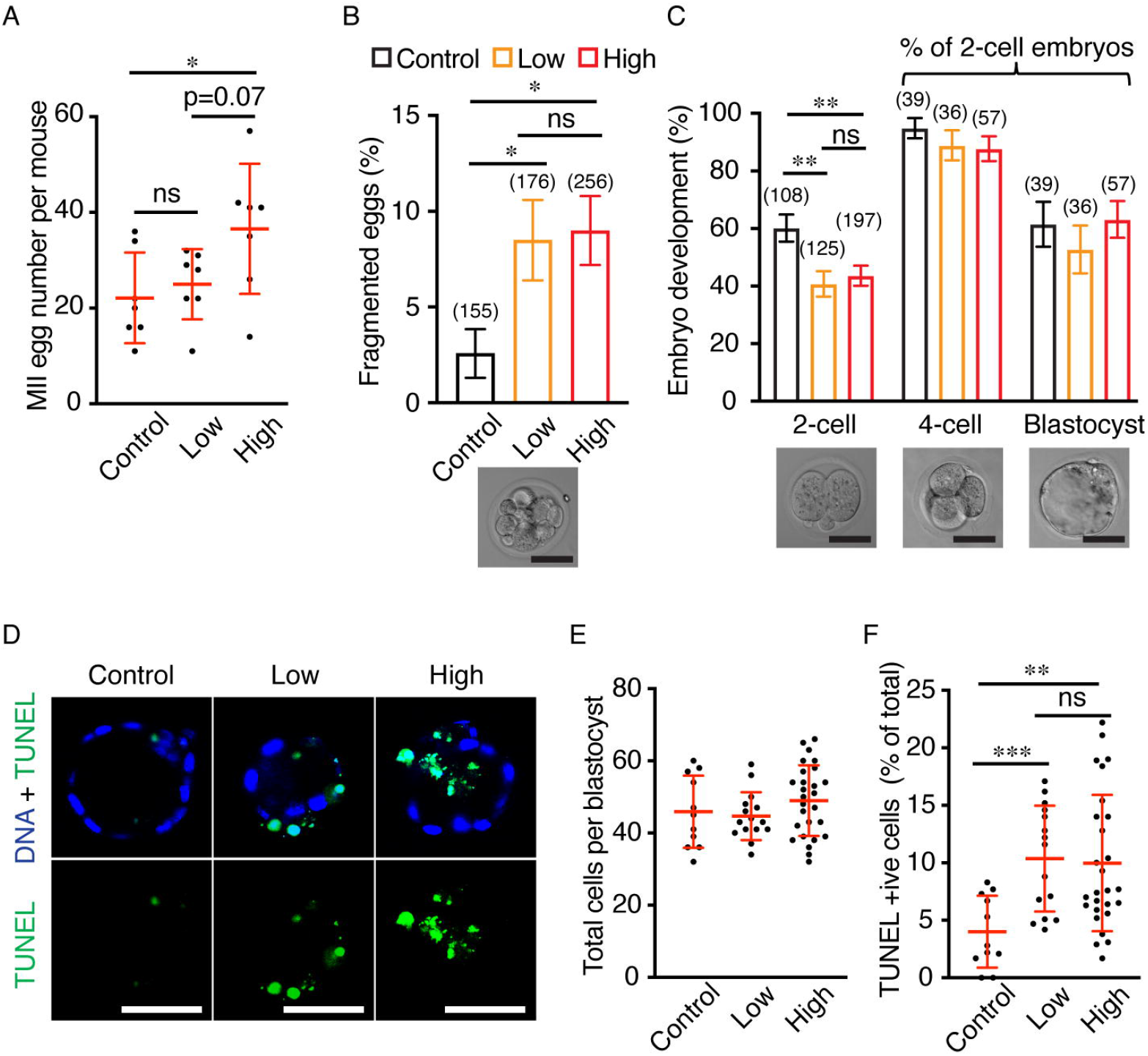
Atrazine exposure during development subsequently impairs *in vitro* fertilization and increases apoptosis in blastocysts. (*A*) Numbers of metaphase II (MII) eggs collected from superovulated females. Low and High refer to the atrazine exposure dose (see text for details). (*B*) Fractions of MII eggs that were fragmented (representative image is shown below; numbers of animals and litters used to generate data in *A* and *B*: control, 7 animals and 4 litters; low dose, 7 and 3; high dose 7 and 3). (*C*) Efficiencies of *in vitro* fertilization (2-cell embryos) and early embryo development (blastocysts; representative images are shown below; efficiencies of 4-cell embryo and blastocyst formation are expressed as percentages of 2-cell embryos; number of animals and litters to generate data for 2-cell embryos are 5 and 3, 6 and 3, 5 and 2 in each exposure group; number of animals and litters to generate data for 4-cell embryos and blastocysts are 3 and 2 in each exposure group). (*D*) Representative images of single *z-*sections of blastocysts derived from *in vitro* fertilization, stained for DNA (Hoechst; blue) and TUNEL (green). (*E*) Total cell numbers per blastocyst were not different between exposure groups. (*F*) Percentage of blastocyst cells that are TUNEL positive cells. Numbers of animals and litters used to generate data in *E* and *F* were 3 and 2, 2 and 2, and 3 and 2 in control, low and high exposure groups, respectively. Numbers of eggs (*B*) or embryos (*C*) analyzed are indicated in parentheses above the bars. Bi-directional error bars represent standard deviation (*A, E, F*) or standard error of a proportion (*B, C*). Data in *A, E*, and *F* were analyzed with unpaired *t* tests; *B* and *C* were analyzed with Fisher’s exact tests. Scale bars in *B-D* represent 50 µm. * indicates *p* < 0.05; **, *p* < 0.01; ***, *p* < 0.001; ns, not significant.

**Figure 3.**
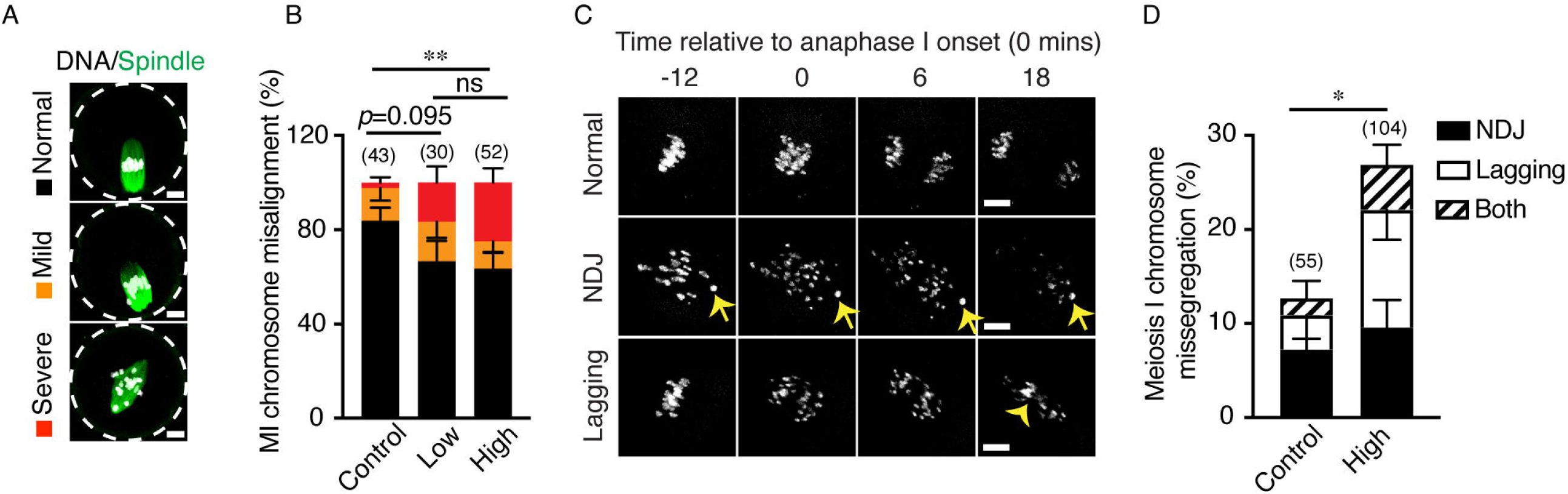
Atrazine exposure during development subsequently causes chromosome misalignment and missegregation in meiosis-I oocytes. (*A*) Representative images of metaphase-I (MI) oocytes, stained for spindles (α-tubulin, green) and DNA (Hoechst, white), illustrating classes of chromosome misalignment. (*B*) Quantification of chromosome misalignment in the MI oocytes represented in panel *A* (data generated from 5 animals from 3 litters in each exposure group). (*C*) Live-cell images from selected timepoints of meiosis-I stage oocytes showing examples of normal chromosome segregation, non-disjunction (NDJ) and lagging segregation. Arrows highlight a non-disjunction event. The arrowhead highlights a lagging chromosome. (*D*) Quantification of chromosome missegregation in meiosis-I oocytes. Data generated from 7 animals and 5 litters (unexposed control); and 8 animals and 4 litters (high dose atrazine), respectively. Numbers of oocytes examined in *B* and *D* are indicated in parentheses above the bars. Error bars represent standard error of a proportion; uni-directional bars are shown for the individual classes to avoid overlaps. Data in *B* and *D* were analyzed with Fisher’s exact tests. Statistical analysis performed in panel *B* compares the distributions of the three alignment classes shown in *A*. All missegregation types were combined for the statistical analysis in panel *D*. Scale bars in *A* and *C* represent 10 µm. *, *p* < 0.05; **, *p* < 0.01; ns, not significant.

**Figure 4.**
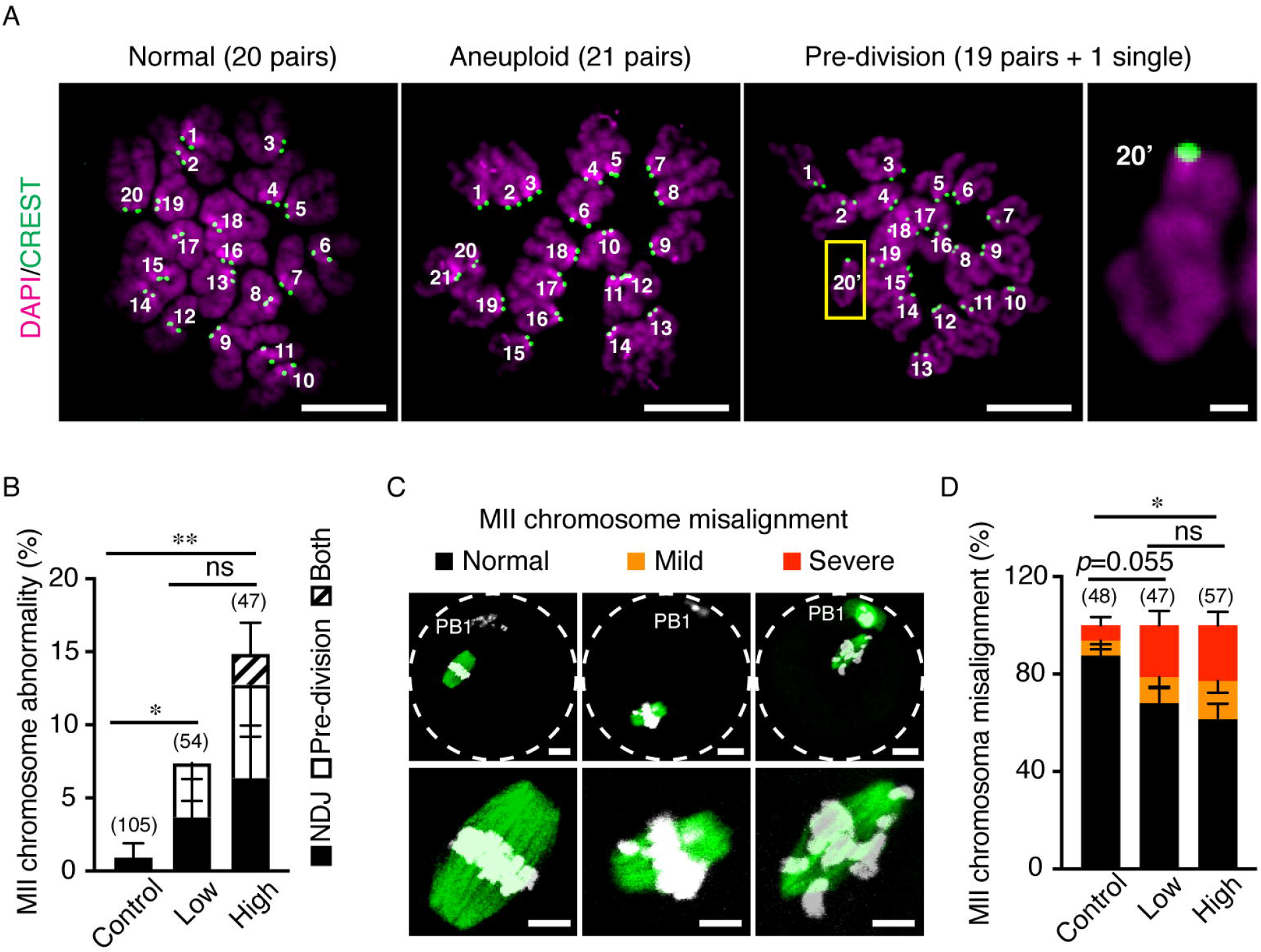
Atrazine exposure during development subsequently causes chromosomal abnormalities and misalignment in metaphase-II eggs. (*A*) Representative images of metaphase-II (MII) oocyte chromosomes showing normal euploid (20 pairs of sister chromatids) and aneuploid (21 pairs) nuclei, and a cell with a single free chromatid (20’; magnified in the right-hand side panels) indicative of a premature separation event (pre-division). Centromeres (green) were immunostained with CREST, and chromosomes (magenta) were counterstained with DAPI. Scale bars represent 10 µm (main panels) and 1 µm (magnified panel). (*B*) Quantification of chromosomal abnormalities in MII eggs from 3-month-old mice (NDJ, nondisjunction). Numbers of animals used were 8 (from 7 litters; unexposed control), 4 (from 3 litters; low dose) and 5 (from 3 litters; high dose), respectively. (*C*) Representative images of MII eggs, stained for spindles (α-tubulin, green) and DNA (Hoechst, white), illustrating classes of chromosome misalignment. PB1, indicates the position of the first polar body. Scale bars represent 10 µm (top panels) and 5 µm (lower panels). (*D*) Quantification of chromosome misalignment in the metaphase-II eggs represented in (*C*). 3 animals were used for each exposure group, from 3 (control), 3 (low), and 2 (high) litters, respectively. Numbers of eggs examined in *B* and *D* are indicated in parentheses above the bars. Error bars represent standard error of a proportion; uni-directional bars are shown for the individual classes to avoid overlaps. Data in *B* and *D* were analyzed with Fisher’s exact tests. All missegregation types were combined for the statistical analysis in panel *B*. Statistical analysis performed in panel *D* compares the distributions of the three alignment classes shown in *C*. * *p* < 0.05; **, *p* < 0.01; ns, not significant.

**Figure 5.**
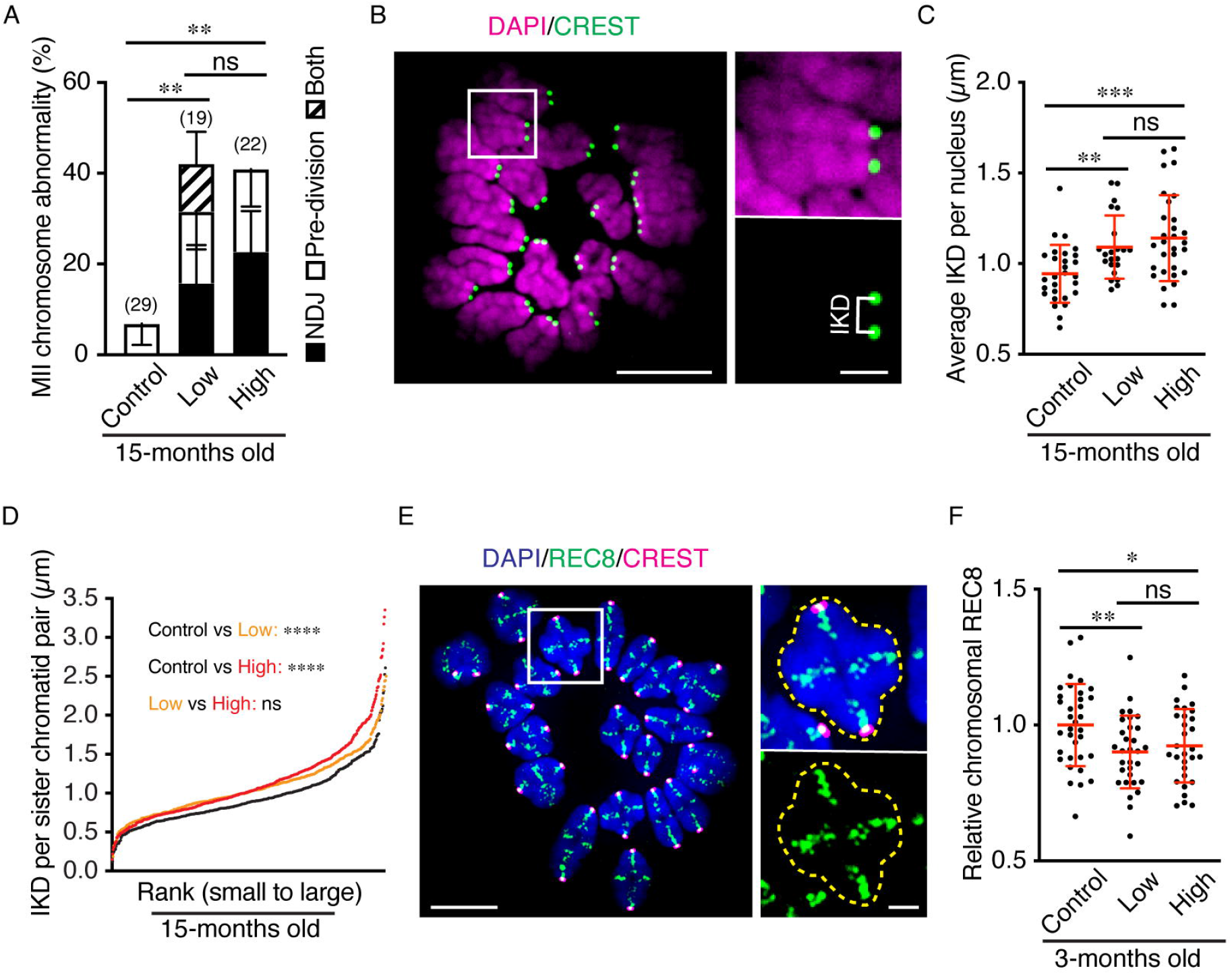
Atrazine exposure in development exacerbates the effects of maternal-age on oocyte quality and weakens sister-chromatid cohesion. (*A*) Quantification of chromosomal abnormalities in metaphase-II (MII) eggs from 15-month-old mice (NDJ, nondisjunction). Numbers of animals used were 8 (from 4 litters; unexposed control), 6 (from 2 litters; low dose), and 5 (from 2 litters; high dose), respectively. (*B*) Representative image of metaphase II chromosomes illustrating the measurement of inter-kinetochore distance (IKD) for a single chromatid pair (magnified panels). Kinetochores (green) were immunostained with CREST, and chromosomes (magenta) were counterstained with DAPI. Scale bars represent 10 µm (main panel) and 2 µm (magnified panels). (*C*) Average IKDs per nucleus for MII eggs from 15-month-old mice. (*D*) Rank distributions of IKDs for individual sister-chromatid pairs of MII eggs from 15-month-old mice. Numbers of sister-chromatid pairs examined in each group were 546 (28 cells, unexposed control), 422 (22 cells, low dose), and 575 (30 cells, high dose), respectively. Numbers of animals used in *C* and *D* were 8 (from 4 litters), 6 (from 2 litters), and 5 (from 2 litters) in control, low, and high dose groups, respectively. (*E*) Representative image of metaphase-I oocyte chromosomes, immunostained for meiosis-specific cohesin component REC8 (green), centromere marker CREST (magenta) and counterstained with DAPI (blue). The right-hand side panels show a single homolog pair; the dashed line highlights the area in which fluorescent intensity was measured to quantify chromosome-associated REC8 (details in the Methods section). Scale bars represent 10 µm (main panel) and 2 µm (magnified panel). (*F*) Quantification of average chromosomal REC8 level per MI oocyte nucleus from 3-month-old mice. Numbers of animals used were 4 (from 3 litters; unexposed control), 4 (from 3 litters; low dose), and 5 (from 3 litters; high dose), respectively. Numbers of eggs examined in *A* are indicated in parentheses above the bars; error bars in *A* show standard error of a proportion; uni-directional bars are shown to avoid overlaps. Bi-directional error bars in *C* and *F* indicate standard deviation. Data were analyzed with Fisher’s exact tests (*A*), unpaired *t* tests (*C, F*), and Mann-Whitney tests (*D*), respectively. MII chromosomal abnormalities were combined for statistical analysis in (*A*). * indicates *p* < 0.05; **, *p* < 0.01; ***, p<0.001; ****, p<0.0001; ns, not significant.

### Confocal Time Lapse Imaging

GV-stage oocytes were cultured *in vitro* for 2 hours, when majority of them had completed GVBD. Oocytes with GVBD were then incubated with 100 nM SiR-DNA (Cytoskeleton, CY-SC007) in M2 medium to label chromosomes until imaging. 4-dimensional imaging was performed using a Zeiss Airyscan LSM 800 confocal equipped with a 40X water immersion objective and Axiocam camera, or a 3i Spinning Disk confocal equipped with a 63X oil immersion objective lens and Flash 4.0 sCMOS camera. Both were equipped with a 37°C temperature-controlled environment. Chromosomes were imaged with 18 z-sections and 2.0 μm z-resolution. The imaging time between consecutive frames was 6 minutes. To capture the events of chromosome alignment and segregation, imaging was performed between 9 and 12 hours of culture, and SiR-DNA containing M2 medium was used throughout the imaging procedure.

### Statistical Analysis

For all comparisons, a minimum of two independent experiments were performed. For each developmental exposure regiment (unexposed control, low and high dose atrazine), 3-16 females from 2-7 different litters were examined, unless otherwise stated. The numbers of mice and litters used for each experiment are stated in the figure legends.

An intrinsic limitation to statistical analysis of oocyte phenotypes is the small numbers of analyzable eggs that can be obtained from individual females. Therefore, oocyte samples from independent experiments are typically pooled to calculate proportions; and comparisons made via Fisher’s exact tests (Maier et al. 2021; Mihajlović et al. 2021). Error bars for analysis of such data indicate the standard error of a proportion (Gruhn et al. 2019). All other mean analyses were performed using either Student’s *t* or Mann– Whitney tests and presented as mean ± SD. Data were processed using GraphPad Prism 8, with a significance threshold of *p* < 0.05. All statistical tests and significance levels are reported in the figure legends.

## RESULTS

To better mimic environmental exposure, female mice were exposed to atrazine in drinking water at two different doses: “high” was a concentration of 33 mg/L, the aqueous saturation level (Yalkowsky et al. 2019) and thus the maximum that will be encountered in contaminated water. Given an average water intake of 5 mL per day and average body weight of 20 g for an adult female, 33 mg/L is equivalent to a dose of ∼8.25 mg/kg/day. For “low” dose exposure, atrazine concentration was 100 µg/L for a dose of ∼25 µg/kg/day. The lower concentration is similar to recorded surface and ground water contamination at some sites (Hatfield et al. 1996; Murphy et al. 2006) and was set as the safe limit in drinking water by the World Health Organization. In rats, prenatal exposure to high dose atrazine (50-100 mg/kg/day) has been associated with pregnancy loss (Narotsky et al. 2001), pup mortality, and reduced bodyweights of offspring (Davis et al. 2011). The two doses employed here did not impact body weight, gestation time, litter size, or pup mortality (Figure S1), indicating that our exposure regimen does not cause gross toxic effects.

### Atrazine Exposure During Development: Meiotic Prophase-I, Fertilization and Early Embryogenesis

The early events of female meiosis and oogenesis are susceptible to perturbations that can reduce the size and quality of ovarian reserves and thereby impact lifetime reproductive success. Key events of meiotic prophase I, including homolog pairing, synapsis and crossing over, occur *in utero*; then around birth, meiosis arrests and oocytes assemble into primordial follicles to establish initial ovarian reserves (Hunter 2017). To determine the effects of atrazine on these critical early events, animals were exposed throughout gestation and early post-natal development by supplying dams with contaminated drinking water while pregnant and nursing, until pups were weaned (Figure 1A). This “developmental” exposure regimen was terminated at embryonic day 18.5 (E18.5), for the analysis of prophase-I events in fetal ovaries; at 18 days post-partum (dpp) for the quantification of ovarian reserves; or at weaning (21 dpp) for subsequent analysis of oocyte quality.

Prophase-I nuclei with one or more defects of chromosome synapsis, including asynapsis and non-homologous pairing of chromosome axes (Figure 1B), were increased ∼3-fold in oocytes from atrazine exposed fetuses (from 5.4% of cells in unexposed controls, to 16.5% and 15.2% in low and high exposure groups, respectively; *p* < 0.01; Figure 1C). Despite this increase in synaptic irregularities, crossover numbers, quantified using the crossover specific marker MLH1, were not significantly changed (Figure 1D and 1E). Comparison of the distributions of MLH1 foci suggests a slight increase (6.9%) in chromosomes with a single crossover at the expense of chromosomes with ≥2 crossovers (7.9% decrease), but only at the higher atrazine dose (Figure 1F; *p* = 0.0125, compared to unexposed controls). However, the frequency of chromosomes lacking an MLH1 focus, indicative of crossover failure, was unchanged.

Prophase-I defects, such as asynapsis, can lead to oocyte apoptosis and thus reduced numbers of primordial follicles (Hunter 2017). However, ovarian reserves (follicle number per ovary from 18 dpp mice) were indistinguishable from unexposed controls (Figure 1G) indicating that oocyte survival is not significantly impacted by the elevated synaptic errors seen in oocytes from atrazine exposed females (Figure 1B and 1C). Atrazine exposure was associated with a slight increase in the number of follicles containing more than one oocyte suggesting that the breakdown of germline cysts (the clusters of oocytes that precede follicle formation)(Pepling 2012) is perturbed by atrazine exposure (Figure 1H and 1I).

The quality of oocytes following developmental atrazine exposure was assessed by measuring the efficiency of *in vitro* fertilization (IVF) and early embryogenesis (2 and 4-cell embryos and blastocyst formation; Figure 2). Metaphase-II (MII) arrested eggs were obtained for IVF from super-ovulated females aged 3-6 months, i.e. 10-23 weeks after atrazine exposure was terminated at weaning. Surprisingly, the number of eggs retrieved from animals exposed to a high dose of atrazine was significantly increased by ∼1.7-fold relative to unexposed controls (Figure 2A, *p* = 0.0399, unpaired *t* test). This observation points to hyperstimulation of follicle growth and resumption of meiosis, or perhaps reduced apoptosis (atresia) of growing follicles. Both low and high dose atrazine-exposure was associated with elevated levels of dead or dying MII eggs with fragmented cellular contents. Only 2.6% of eggs from unexposed females were fragmented compared to 8.5% and 9.0% from low and high dose exposure cohorts, respectively (*p* = 0.0307 and 0.0125 respectively, Fisher’s exact test; Figure 2B).

To determine the developmental capacity of oocytes, intact MII eggs from exposed and control animals were fertilized with sperm from unexposed males. 60.2% of eggs from unexposed females formed 2-cell embryos 24 hrs after eggs and sperm were mixed (Figure 2C). IVF efficiency dropped by around one third for eggs from atrazine-exposed animals, down to 40.8% and 43.7% 2-cell embryos from the low and high dose groups, respectively (*p* = 0.0038 and 0.0061 respectively, Fisher’s exact test). In embryos that successfully traversed the 2-cell stage, the efficiency of 4-cell embryo and blastocyst formation was indistinguishable between exposed and unexposed groups (Figure 2C). Blastocyst integrity was assessed by counting total cell number, and the percentage of cells that were apoptotic (Figure 2D–F). Total cell number did not differ for blastocysts derived from atrazine exposed and control groups (Figure 2E). However, both low and high dose atrazine exposure were associated with ∼2-fold increases in TUNEL-positive apoptotic cells (10.4 ± 4.6% and 10.0 ± 5.9% in low and high dose exposures versus 4.0 ± 3.1% in unexposed controls, mean ± S.D; *p* = 0.0006 and 0.0033 respectively, unpaired *t* test; Figure 2F). Moreover, in blastocysts derived from eggs of atrazine-exposed animals, the fraction of cells that were apoptotic was very variable, ranging from 1.7 to 22.2%. Blastocysts with a high fraction of apoptotic cells could be a consequence of problems present in the zygote or very early embryo such as aneuploidy, which does not necessarily impede blastocyst formation but leads to increased apoptosis (Bolton et al. 2016; Rubio et al. 2007; Tšuiko et al. 2019). The reason for the reduced fertilization efficiency following atrazine exposure is less clear. Possibly, atrazine causes epigenetic changes that perturb the transcriptional responses required for optimal maturation and fertilization (Gely-Pernot et al. 2017). Together, our analysis reveals that atrazine exposure during development has long-term negative impacts on various indicators of oocyte quality and results in higher levels of fragmented MII eggs, lower fertilization rates, and increased apoptosis in blastocysts.

### Chromosome Misalignment and Missegregation in Oocytes From Atrazine Exposed Animals

To better understand the etiology of reduced egg quality following atrazine exposure during development, we cultured germinal vesicle (GV) stage oocytes and analyzed chromosome integrity and segregation at meiosis I. Atrazine exposure did not perturb maturation, with very similar efficiencies of meiotic resumption and completion of meiosis I observed for oocytes from all three exposure groups (assayed respectively via germinal vesicle breakdown (GVBD) and first polar body extrusion (PBE); Figure S2A, B and C).

#### Metaphase-I Abnormalities

Chromosome congression is particularly important for accurate segregation in mammalian oocytes, since the spindle assembly checkpoint is less robust in meiosis, such that misaligned chromosomes can escape and missegregate (Lane et al. 2012; Nagaoka et al. 2011). Congression at metaphase I was analyzed in fixed oocytes fluorescently stained to visualize spindles and chromosomes (Figure 3A). Defects were classified as mild, with 1 or 2 misaligned chromosomes, or severe with ≥3 misaligned chromosomes (Figure 3A). 16.3% of control oocytes showed chromosome misalignment; 14.0% were classified as mild and only 2.3% as severe (Figure 3B). In low and high dose atrazine groups, misalignment was seen in 33.4% and 36.5% of oocytes, respectively, with 16.7% and 25.0% showing severe misalignment. Statistical analysis of the distributions of the three alignment classes showed a significant increase in chromosome misalignment for the high-dose group (*p* = 0.0050, Fisher’s exact test) but not for the low dose group (*p* = 0.0950). Increased misalignment in metaphase I predicts chromosome missegregation at anaphase I. To test this inference, live-cell imaging was used to quantify meiosis-I chromosome missegregation, i.e. non-disjunction and lagging chromosomes (Figure 3C; Movies S1-S3). Missegregation was detected in 12.7% of control oocytes (Figure 3D), a higher frequency than typically observed in the literature (≤5%, (Chiang et al. 2010; Yun et al. 2014b)) that appeared to be caused by the SiR-DNA fluorogenic DNA label (Spirochrome) used to visualize chromosomes.

Notwithstanding this higher baseline, oocytes from animals exposed to the higher dose of atrazine had more than twice as many chromosome segregation errors as unexposed controls (26.9%, *p* = 0.0453, Fisher’s exact test). Thus, consonant with our analysis of fixed cells, atrazine exposure during development led to elevated levels of chromosome misalignment and missegregation at meiosis I.

#### Metaphase-II Abnormalities

Chromosome misalignment and missegregation during meiosis I are expected to manifest as chromosomal abnormalities in the succeeding metaphase-II arrested eggs. To test this prediction, oocytes from control animals and animals exposed to atrazine during development were cultured through metaphase-II arrest and chromosome prepared for immunofluorescence imaging. Euploid metaphase-II mouse eggs contain 20 pairs of sister chromatids connected at their centromeric ends, while aneuploid eggs have either extra or missing sister chromatid pairs. A second type of chromosomal abnormality – pre-division or precocious separation of sister chromatids – results in free single chromatids that are expected to segregate randomly during meiosis II (upon fertilization) and are therefore a high risk for embryo aneuploidy (Figure 4A). Consistent with previous studies in the C57Bl/6J mouse strain (Yun et al. 2014a), only 1.0% of control eggs were chromosomally abnormal. Strikingly, this rate increased to 7.4% and 14.9%, respectively, for eggs from the low and high dose atrazine exposure groups (*p* = 0.0458 and 0.0012, respectively, Fisher’s exact test; Figure 4B).

Chromosome alignment in metaphase-II oocytes was also examined. As predicted from the high levels of chromosome abnormalities seen in oocytes from atrazine-exposed animals, ≥30% of metaphase-II oocytes from the low and high dose groups showed misalignment, with severe defects (≥3 chromosomes) being seen in more than 20% of cells (Figure 4C and 4D). Again, comparison of the distributions showed that this effect was only significant in the high dose group (*p* = 0.0551 and 0.0101 for the low and high-dose groups, respectively); although when analyzed as a simple proportion, there were significantly more cells with chromosome misalignment (mild + severe) in the low dose group (*p* = 0.0270). Misalignment in metaphase-II arrested oocytes, even if they are euploid, will elevate the risk of ensuing embryo aneuploidy following fertilization and completion of MII.

We conclude that atrazine exposure during development has long-term negative consequences for oocyte meiosis specifically: (1) chromosome misalignment at both metaphase I and II; and (2) chromosome missegregation and pre-division presenting in metaphase II.

### Exacerbation of the Maternal Age Effect

With our developmental exposure regimen, oocyte defects are observed at 3 months, long after atrazine is withdrawn at the time of weaning (21 dpp; Figures 2 through 4). Therefore, we explored the relationship between exposure to atrazine during development, and the increased chromosome errors seen with advanced maternal age (Chiang et al. 2010; Lister et al. 2010; Yun et al. 2014b). The C57Bl/6J strain employed here experiences a relatively minor age-related increase in meiotic segregation errors, from a frequency of 1.0% of oocytes from 3-month-old animals to 6.9% at 15-months of age (Figure 5A). Both low and high dose atrazine exposure dramatically exacerbated age-related errors, resulting in chromosomal abnormalities in ≥40% of metaphase II eggs from 15-month-old animals (Figure 5A). Thus, atrazine exposure during development acted synergistically with maternal age, causing very high levels of chromosomally abnormal oocytes in older females.

Progressive depletion of sister-chromatid cohesion, leading to chiasma loss (and the appearance of univalent chromosomes) and premature separation of sister chromatids, is a major cause of the high levels of chromosomal errors seen in oocytes from older females (Sakakibara et al. 2015; Zielinska et al. 2015). Therefore, we assessed whether cohesion is weakened following atrazine exposure during development. Increased inter-kinetochore distance (IKD) is a readout of weakened cohesion between sister centromeres (Figure 5B; (Chiang et al. 2010; Lagirand-Cantaloube et al. 2017)). Average IKDs were significantly increased in metaphase-II nuclei from 15-month-old females following atrazine exposure during development; from 0.94 ± 0.16 µm in unexposed controls to 1.09 ± 0.17 µm and 1.14 ± 0.24 µm in low and high dose atrazine groups, respectively (Figure 5C; *p* = 0.0032 and *p* = 0.0005, respectively, unpaired t-test). Rank distributions of the IKDs of individual sister-chromatid pairs confirmed that centromeric cohesion is weakened following atrazine exposure (Figure 5D; *p* < 0.0001, for both low and high dose groups, Mann-Whitney test).

Weakening of cohesion following atrazine exposure was further analyzed by quantifying levels of the cohesin complexes that are responsible for sister-chromatid cohesion (Cheng and Liu 2017). Metaphase-I oocyte chromosomes from three-month-old females were immunostained for the meiosis-specific cohesin subunit REC8 and fluorescence intensity was calculated (Figure 5E and 5F; staining was performed at 3 months because REC8 is very depleted and difficult to detect in oocytes from very old animals (Chiang et al. 2010)). This analysis revealed reduced levels of chromosome-associated cohesin in metaphase-I oocytes from atrazine exposed animals; relative REC8 was reduced by 9.9% and 7.6% in the low and high dose atrazine groups, respectively (Figure 5F; *p* = 0.0070 and *p* = 0.0382, respectively, unpaired *t* test). Thus, atrazine exposure mimics and thereby exacerbates the effects of maternal aging on oocyte quality by weakening sister-chromatid cohesion.

### Atrazine Exposure During Adulthood and Oocyte Abnormalities

Although atrazine exposure during development (gestation through weaning) clearly impacts oocyte divisions, the critical events that occur during fetal development (chromosome synapsis and crossing over) and the early postnatal period (formation of primordial follicles) were only very mildly perturbed (Figure 1). These observations suggest that the critical exposure window may occur later in life between 4 dpp, when primordial follicles are formed, and weaning at 21 dpp. This possibility was tested by exposing adult mice (at 2-months old) to atrazine-contaminated drinking water for 3 months (at the high and low doses detailed above, (Figure 6A), after which oocyte divisions were analyzed (Figure 6B–F).

**Figure 6.**
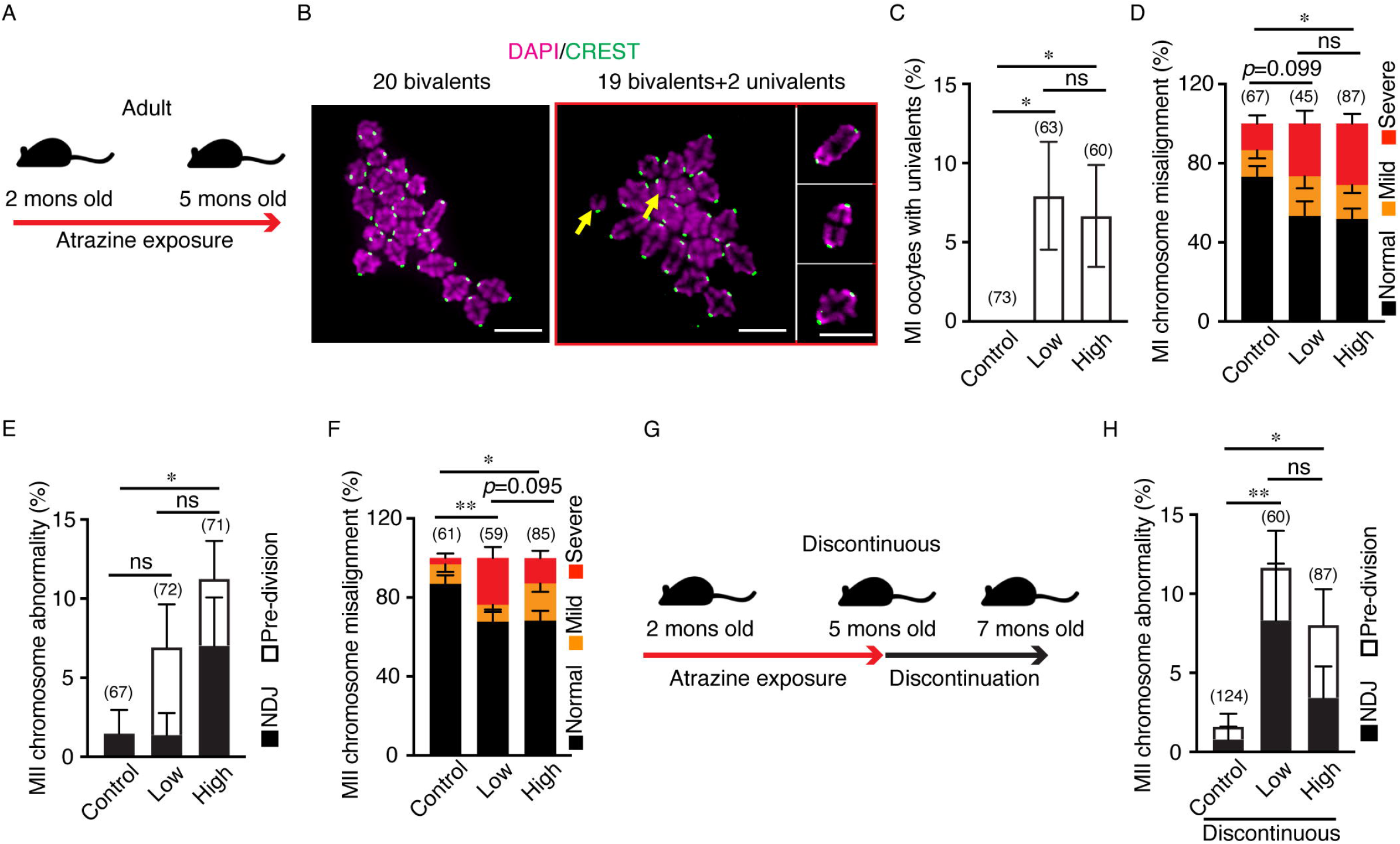
Atrazine exposure during adulthood causes persistent oocyte chromosome defects. (*A*) Schematic of atrazine exposure regimen in adult females. (*B*) Representative images of metaphase-I oocyte chromosome preparations showing a normal nucleus with 20 bivalents and a nucleus containing 19 bivalents and two univalents (yellow arrows). Chromosomes shown in insets are from the same cell, but were in different fields of view. DNA is colored magenta and centromeres are green. Scale bars represent 10 µm. (*C*) Quantification of unconnected univalent chromosomes in chromosome preparations from metaphase-I oocytes. Numbers of animals used were 6 (unexposed control), 5 (low dose), and 5 (high dose), respectively. (*D*) Quantification of chromosome misalignment in metaphase-I oocytes. Numbers of animals used were 4, 3, and 4, respectively. (*E*) Quantification of chromosomal abnormalities in metaphase-II eggs. Numbers of animals used were 7, 6, and 6, respectively. (*F*) Quantification of chromosome misalignment in metaphase-II eggs. Numbers of animals used were 4, 3, and 4, respectively. (*G*) Schematic of the discontinuous atrazine exposure regimen. (*H*) Quantification of chromosomal abnormalities in metaphase-II eggs following discontinuous atrazine exposure. Numbers of animals used were 9, 4, and 8, respectively. Numbers of MI oocytes (*C, D*) and MII eggs (*E, F, H*) examined are indicated in parentheses above the bars. Error bars represent standard error of a proportion; uni-directional bars are shown in *D–F*, and *H* to avoid overlaps. Data were analyzed with Fisher’s exact tests. Statistical analysis performed in *D* and *F* compares the distributions of the three alignment classes. MII chromosomal abnormalities were combined for statistical analyses in panels *E* and *H*. * indicates *p* < 0.05; **, *p* < 0.01; ns, not significant.

As observed for oocytes from females exposed during development, neither low nor high dose atrazine exposure during adulthood affected the maturation efficiency of oocytes when cultured *in vitro*. Specifically, oocytes from all examined groups completed GVBD and PBE with efficiencies of ≥75% (Figure S2D and E). However, analysis of metaphase-I chromosome preparations revealed increased levels of univalent chromosomes in oocytes from atrazine-exposed animals (Figure 6B and 6C). Univalents were never observed in controls but were present in 7.9 % and 6.7 % of oocytes from the low and high dose atrazine groups, respectively (*p* = 0.0195 and 0.0391 respectively, Fisher’s exact test). Univalent formation is consistent with a weakening of sister-chromatid cohesion following atrazine exposure (Figure 5E and 5F).

The presence of univalents during metaphase I anticipates defective congression and segregation (Nagaoka et al. 2011). Indeed, nearly 50% of oocytes from both low and high dose atrazine-exposed adults had congression defects at metaphase I (Figure 6D). Comparison of the three alignment classes showed a significant increase in misalignment for the high dose group (*p* = 0.0170, Fisher’s exact test), but not for the low dose group (*p* = 0.0991). However, when analyzed as a simple proportion, cells with chromosome misalignment (mild + severe) were significantly increased in the low dose group (*p* = 0.0429). In the high dose group, severe chromosome misalignment (≥3 chromosomes) was seen in 31.0% of cells compared to 13.4 % of control oocytes. These levels are much higher than expected if univalents were the sole cause of misalignment, implying that atrazine causes additional defects that perturb congression such as merotelic spindle attachment that can result from weakening of cohesion between the centromeres of sister-chromatids (Figure 5B–D)(Shomper et al. 2014; Zielinska et al. 2015).

In metaphase-II arrested oocytes, chromosomal abnormalities were observed in 6.9% and 11.3% of oocytes from the low and high dose groups, respectively, compared to only 1.5% in controls (Figure 6E; again, only the high-dose group was significantly different to controls, *p* = 0.0337, Fisher’s exact test). Chromosome congression was also analyzed in metaphase-II arrested eggs, revealing misalignment in over 30% of eggs from atrazine exposed groups compared to 13.1% in controls (Figure 6F; *p* = 0.0033 and 0.0302 for low and high dose groups respectively, Fisher’s exact test).

In conclusion, atrazine exposure during adulthood causes defects in oocyte divisions that are the same magnitude as those seen following developmental exposure. Although it is formally possible that exposure during development and adulthood cause the same defects through distinct mechanisms, the most straightforward interpretation is that the critical period for atrazine exposure occurs after the perinatal period when ovarian reserves are established. These considerations suggest that established primordial follicles may be a primary target of atrazine perturbation.

### Atrazine Exposure in Adults and Long Term Impacts on Oocyte Quality

To test whether exposure in adulthood has long-term effects, as seen following developmental exposure, mice were given atrazine-contaminated drinking water between 2-5 months of age, followed by pure water for an additional 2 months before MII-arrested oocytes were analyzed (Figure 6G). This analysis confirmed that the effects of atrazine exposure are persistent, with significantly elevated levels of chromosomal abnormalities being observed in both low and high dose groups, that were the same magnitude as those seen following continuous exposure immediately prior to oocyte analysis (Figure 6H; compare with 6E).

## DISCUSSION

Our analysis shows that chronic ingestion of low levels of atrazine via drinking water results in persistent negative effects on female reproduction in mouse, with defects being detected during meiosis, fertilization and early development. Importantly, we provide the first clear evidence that atrazine exposure perturbs chromosome segregation during both meiotic divisions, and dramatically exacerbates the effects of maternal age on oocyte quality. Mechanistically, one cause of these defects appears to be atrazine-induced weakening of sister-chromatid cohesion, effectively mimicking a well-characterized facet of oocyte aging (Cheng and Liu 2017; Jessberger 2012). Our data are summarized and synthesized into a model that reconciles the chromosomal perturbations observed following atrazine exposure (Figure 7).

**Figure 7.**
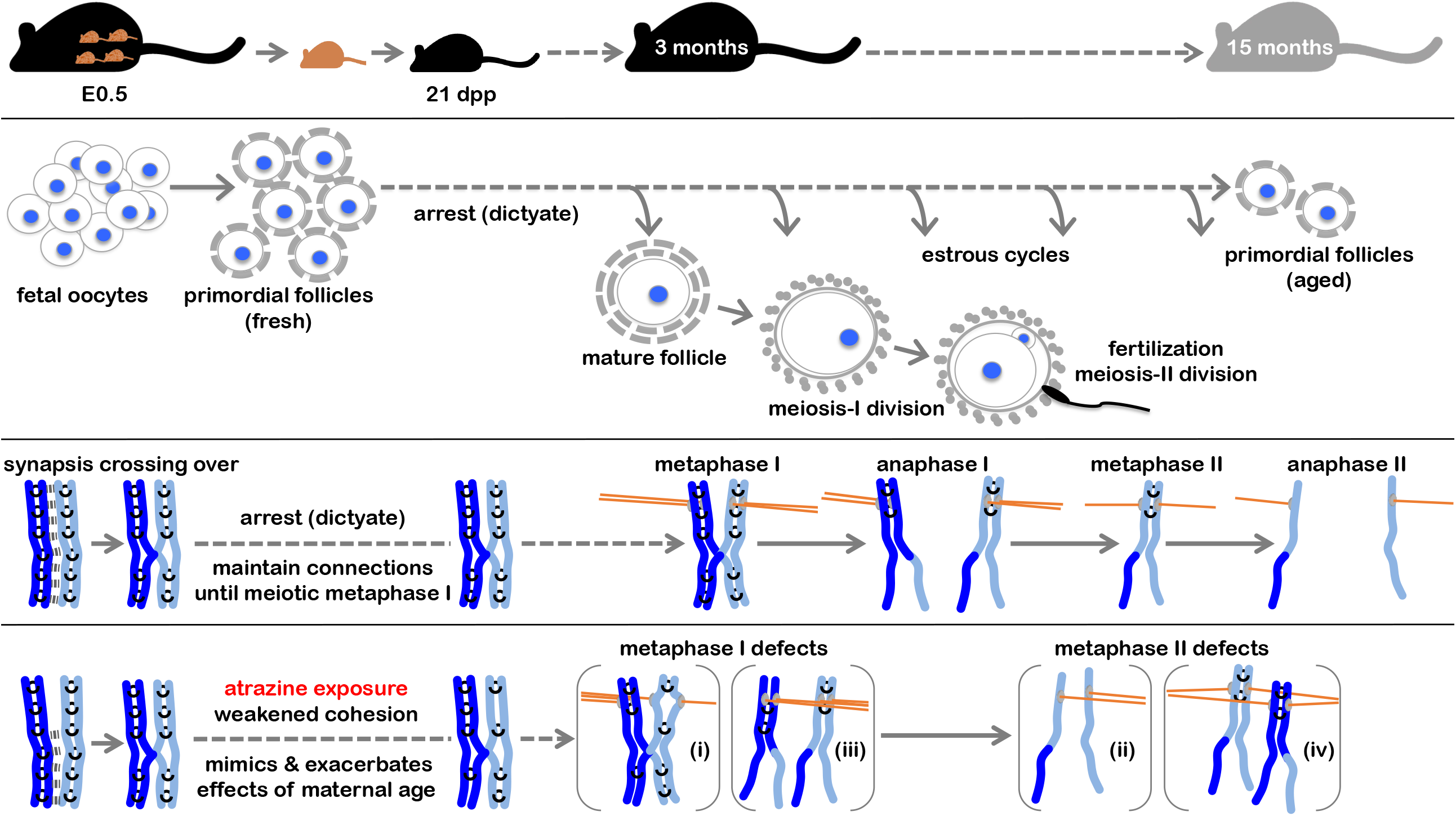
Summary of the long-term effects of atrazine exposure during development on oocyte quality. *First row:* timeline of mouse development and window of atrazine exposure (developmental exposure regimen) and ages at which parameters of oocyte quality were assessed. *Second row:* timeline of oocyte development. The events of meiotic prophase-I occur in fetal oocytes, which then arrest around birth and assemble into primordial follicles. With each estrus cycle, cohorts of follicles grow, and meiosis resumes in dominant follicles that are about to be ovulated. The meiosis-I division ensues, and eggs then arrest at metaphase-II until fertilization triggers division. *Third row:* the chromosomal events of meiosis begin in fetal oocytes with chromosome pairing, synapsis (synaptonemal complex indicated by a thick dashed line) and crossing over, shown here for a single pair of homologs (dark and light blue lines). Each homolog comprises a pair of sister-chromatids connected by cohesins (black rings); consequently, crossing over results in the connection of homologs, which enables their bipolar alignment on the metaphase-I spindle and accurate segregation (disjunction) at meiosis I (grey discs, kinetochores; orange lines, microtubules). Thus, connections must be maintained in primordial follicles throughout their arrest period. Connections are resolved at anaphase-I by the cleavage of cohesins, allowing homologs to separate. Cohesins that connect sister centromeres are protected from cleavage until anaphase-II, allowing accurate congression and segregation of sister chromatids. *Fourth row:* chromosomal errors in oocytes following atrazine exposure. In fetal oocytes, synaptic errors were modestly increased but numbers of crossovers were not significantly changed. We propose that primordial follicles are the consequential target of atrazine exposure, which mimics and exacerbates the effects of maternal age by weakening sister-chromatid cohesion, indicated by fewer cohesin complexes (black rings). Weakening of cohesion between centromeres can result in merotelic attachment (**i**), misalignment at metaphase I, and chromosome lagging at anaphase I. If sister chromatids prematurely separate in anaphase I, the free chromatids are prone to misalign in metaphase II (**ii**) and missegregate in anaphase II leading to aneuploidy. Alternatively, sister chromatids may prematurely separate in metaphase II and missegregate in anaphase II. Loss of cohesion distal to crossover points results in prematurely separated univalents (**iii**), which can co-orient at metaphase I, and co-segregate at anaphase I (nondisjunction); resulting in aneuploidy in metaphase II (**iv**) and in the ovum pronucleus following fertilization. Alternatively, the sister-chromatids of a univalent may undergo premature segregation in anaphase I (reverse segregation).

### Meiotic Prophase I and Follicle Formation

Defects in homolog synapsis were increased following exposure *in utero* (Figure 1C), but the impact on crossing over and oocyte survival was negligible (Figure 1E-G). A previous mouse study detected a slight reduction in crossovers (∼6% fewer MLH1 foci) with acute dosing of pregnant dams (100 mg/kg/day), perhaps as a consequence of the higher atrazine dose employed (12 and 4,000-fold higher than the two doses employed in our study) (Gely-Pernot et al. 2017). Slight increases in multi-oocyte follicles were detected following exposure to ∼8.25 mg/kg/day atrazine (this study, Figure 1H and 1I) and to the much higher dose of 100 mg/kg/day (Gely-Pernot et al. 2017). This defects could be explained by documented increases in progesterone (Goldman et al. 2013; Graceli et al. 2020; Wirbisky and Freeman 2015), shown by Chen et al. (2007) to impede oocyte-nest breakdown and the assembly of primordial follicles in mouse ovaries (Chen et al. 2007). Moreover, the enhanced ovulation that we observed following atrazine exposure (Figure 2A) could also be explained by elevated progesterone, since in an *ex-vivo* ovary culture model, 100 ng/ml progesterone was able to promote initial follicle growth and increase the numbers of ovulated oocytes (although higher concentration, 1 µg/ml, inhibited ovulation) (Komatsu and Masubuchi 2017).

### Weakened Cohesion Explains Chromosomal Abnormalities Following Atrazine Exposure

Since exposure during development or adulthood causes similar levels of chromosome errors, we favor the idea that quiescent oocytes present in primordial follicles are the consequential targets of atrazine (Figure 7, 2^nd^ row). Efficient chromosome segregation requires that primordial follicles maintain a critical level of sister-chromatid cohesion until they mature and complete meiotic divisions (Greaney et al. 2018). Optimal cohesion has two critical functions for chromosome alignment and segregation during the two divisions of meiosis:

i. cohesion between centromeres helps organize the kinetochores of sister chromatids into a fused structure required for monopolar attachment and thus correct orientation on the metaphase I spindle (Figure 7, 3^rd^ row) (Greaney et al. 2018; Wang et al. 2019). Suboptimal centromere cohesion causes premature individualization of kinetochores allowing bipolar attachment of sister chromatids to the metaphase I spindle (merotelic attachment, abnormality (i) in Figure 7, 4^th^ row). Premature segregation of sister-chromatids may then ensure at anaphase I, with a high probability of producing an aneuploid ovum pronucleus after anaphase II. Even without erroneous segregation of sister-chromatids at anaphase I, weakening of centromere cohesion may lead to premature separation and coorientation of sister chromatids in metaphase II (abnormality (ii) in Figure 7, 4^th^ row), also a high risk for aneuploidy.
ii. cohesion between chromosome arms maintains connections (chiasmata) between homolog pairs that enable their stable bipolar orientation and congression on the metaphase-I spindle (Figure 7, 3^rd^ row). Suboptimal arm cohesion can lead to premature loss of inter-homolog connections, i.e. univalent formation. Univalents may coorientate on the metaphase-I spindle (abnormality (iii) Figure 7, 4^th^ row) and cosegregate (nondisjunction) at anaphase I. The resulting aneuploidy at metaphase II (abnormality (iv) Figure 7, 4^th^ row), will result in disomy in the ovum pronucleus. Alternatively, the sister-chromatids of a univalent may biorient in metaphase I and segregate in anaphase I (“reverse segregation”) (Ottolini et al. 2015; Sakakibara et al. 2015).

Thus, the weakening of sister-chromatid cohesion caused by atrazine exposure can explain the elevated levels of chromosome misalignment and abnormalities seen in both metaphase I and II oocytes.

### Exacerbation of Maternal Age Effect

Weakened sister-chromatid cohesion can also explain how atrazine exposure exacerbates the effects of maternal age on oocyte quality. Egg aneuploidy increases with advancing maternal age (Cheng and Liu 2017; Jessberger 2012). Specifically, a physiological aging process associated with ovulation is inferred to cause progressive loss of cohesion in primoradial follicles (Chatzidaki et al. 2021). When residual cohesion falls below a critical threshold (Chiang et al. 2010; Jessberger 2012), chromosomal errors increased dramatically. Thus, with respect to weakening cohesion, atrazine exposure appears to mimic maternal aging, accelerating the age at which cohesion is reduced below the critical threshold required for accurate segregation.

### Mechanism of Action

How atrazine exposure causes the meiotic aberrations and cohesion weakening observed here remains unclear. Across vertebrate classes, atrazine acts through multiple mechanisms to disrupt the neuroendocrine system, especially steroidogenesis pathways (Goldman et al. 2013; Graceli et al. 2020; Hayes et al. 2011; Wirbisky and Freeman 2015). Atrazine may also impact oocyte function through documented increases in oxidative stress including mitochondrial dysfunction seen in both aquatic organisms and mammals (Gely-Pernot et al. 2017; Semren et al. 2018). Moreover, fetal exposure to high-levels of atrazine caused reduced expression of genes involved in protection from oxidative stress in the ovaries of 6-day-old mice (Gely-Pernot et al. 2017). It will be important to establish whether similar effects on the oxidative stress response are caused by low-dose atrazine exposure at different times during development, and in adults. Importantly, oxidative damage is suggested to be a possible cause of cohesion loss as primordial oocytes age (Perkins et al. 2016; Sasaki et al. 2019; Semren et al. 2018) and is therefore a good candidate to explain how atrazine exposure interacts with maternal age to reduce oocyte quality. Also, given that ovulation frequency appears to be the physiological basis of oocyte aging (Chatzidaki et al. 2021), the enhanced ovulation observed following atrazine exposure could accelerate this process.

Extensive similarities between mouse and human oocytes suggest that our findings may be pertinent for the reproductive health of human females; and set the stage for expanded studies in rodents, non-human primates and humans that will inform and guide the public, health workers and policymakers.

## Acknowledgments

We thank Richard Schultz and members of the Hunter Lab for support and discussions; and Scott Keeney for the kind gift of REC8 antibodies. Y.Y. and N.H. designed the research; Y.Y., C.S., R.M. and M.H. performed the experiments; Y.Y., S.L., C.S., R.M., C.K, T.W. and N.H. analyzed data; Y.Y. and N.H. wrote the paper. This research was supported by the National Institute of Environmental Health Sciences under Award P30ES023513. N.H. is an Investigator of the Howard Hughes Medical Institute, which also supported this study.

