## Supplemental Material for "Reduced oocyte quality and exacerbation of the maternal age effect are enduring consequences of low-level atrazine exposure in mouse"

**This PDF file includes:**

Figures S1 to S2

Legends for Movies S1 to S3

**Other supplementary materials for this manuscript include the following:**

Movies S1 to S3

**Figure S1**

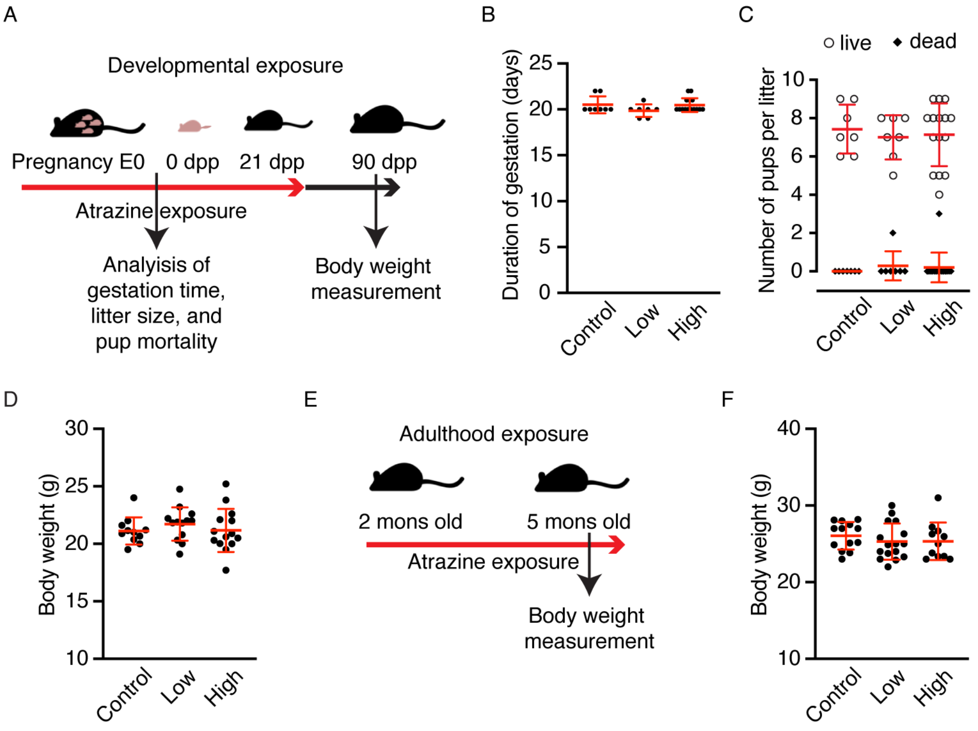


**Figure S1.** Body weight, gestation time, litter size, and pup mortality are not altered by the atrazine doses employed in this study. (*A*) Schematic of developmental exposure regimen indicating times of measurements (E0, embryonic day zero; dpp, days post-partum). (*B*) Gestation time was not significantly altered by exposure to atrazine. Number of litters analyzed were 8, 7, and 15, respectively for unexposed controls, low dose and high dose groups, respectively. (*C*) Number of live (open circles) and dead (filled diamonds) pups delivered per litter were not altered following atrazine exposure. Number of litters analyzed were 7, 7, and 15 in unexposed controls, low and high dose groups, respectively. (*D*) Body weights at 3-months old following developmental atrazine exposure. Number of animals weighed were 11 (from 3 litters), 13 (from 3 litters), and 14 (from 5 litters) for unexposed controls, low dose and high dose groups, respectively. (*E*) Schematic of atrazine exposure regimen in adult females. Body weights were measured at 5-months old. (*F*) Body weights at 5-months old following atrazine exposure during adulthood. Number of animals weighed were 13, 16, and 11, for unexposed controls, low dose and high dose groups, respectively. Bi-directional error bars represent standard deviation. Data were analyzed with either Mann-Whitney test (*B*) or unpaired *t* tests (*C, D* and *F*).

**Figure S2**

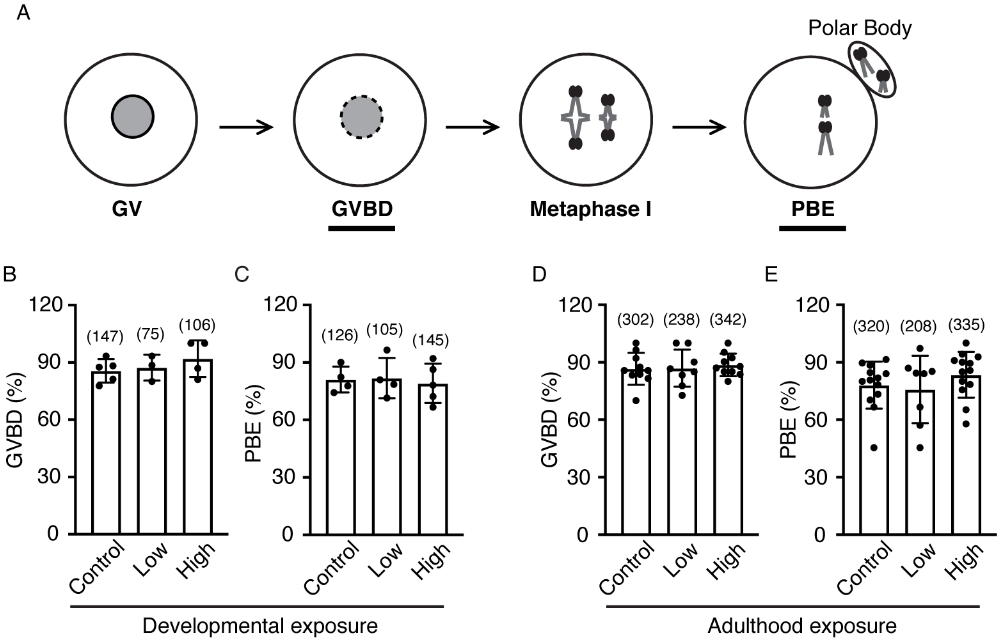


**Figure S2.** Analysis of oocyte maturation *in vitro* following atrazine exposure during development or adulthood. (*A*) Cartoon illustrating the processes analyzed in panels *B–E.* Germinal vesicle breakdown (GVBD) is a readout of the resumption of meiosis; and first polar body extrusion (PBE) indicates completion of the meiosis-I division. (*B* and *D*) Efficiency of GVBD in oocytes harvested from unexposed controls and females exposed to low and high doses of atrazine during development (*B*) or adulthood (*D*), respectively. (*C* and *E*) Efficiency of PBE in oocytes harvested from unexposed controls and females exposed to low and high doses of atrazine during development (*C*) or adulthood (*E*), respectively. No significant differences were detected between exposed and control groups. Numbers of oocytes analyzed are indicated in parentheses. Bi-directional error bars represent standard deviation. Data were analyzed using unpaired *t* tests. Numbers of animals used for control, low and high dose groups were 5 (from 3 litters), 3 (from 2 litters), and 4 (from 2 litters) in (*B*); 4, 4, and 5 in (*C;* all from 3 litters); 10, 8 and 10 in (*D*); and 13, 8 and 13 in (*E*).

**Movie S1. Live-cell imaging of a mouse oocyte showing normal chromosome segregation in meiosis I.** Chromosomes were labeled with SiR-DNA. Elapsed time is shown relative to the onset of anaphase I (hh:mm). Scale bar represents 10 μm.

**Movie S2. Live-cell imaging of a mouse oocyte showing homolog nondisjunction in meiosis I.** Chromosomes were labeled with SiR-DNA. Elapsed time is shown relative to the onset of anaphase I (hh:mm). Scale bar represents 10 μm.

**Movie S3. Live-cell imaging of a mouse oocyte showing chromosome lagging at anaphase I.** Chromosomes were labeled with SiR-DNA. Elapsed time is shown relative to the onset of anaphase I (hh:mm). Scale bar represents 10 μm.
